# Identification of GTF2IRD1 as a novel transcription factor essential for acute myeloid leukemia

**DOI:** 10.1101/2022.08.09.503300

**Authors:** Yaser Heshmati, Gözde Türköz, Joanna E. Zawacka, Marios Dimitriou, Aditya Harisankar, Johan Boström, Huan Cai, Nadir Kadri, Mikael Altun, Hong Qian, Julian Walfridsson

## Abstract

Acute myeloid leukemia (AML) is an aggressive blood malignancy characterized by clonal accumulation of immature myeloid progenitors in the bone marrow and peripheral blood. Transcription factors are the most frequently mutated and dysregulated genes in AML, and they have critical roles in AML pathogenesis and progression. In this study, we performed large-scale RNA interference screens in *MLL-AF9*-transformed AML cells and identified *GTF2IRD1* as a novel transcription factor essential for the survival of various types of myeloid leukemic cells *in vitro* and *in vivo*, but not for primary normal hematopoietic cells. Inhibition of *GTF2IRD1* reduced the frequency of primary childhood and adult AML cells, including cell populations enriched for leukemia-initiating cells. In animal models for AML, inhibition of *GTF2IRD1* significantly delayed the disease progression. Loss of *GTF2IRD1* promoted accumulation of AML cells in the G0 phase of the cell cycle but caused minor effects in apoptosis induction. In line with this, RNA sequencing analysis revealed a significant downregulation of E2F targets as a consequence of genetic inhibition of GTF2IRD1. Taken together, we identified GTF2IRD1 as a transcription factor with a selective importance in AML, and our findings may contribute to the development of improved therapeutic inventions for the disease.

## Introduction

Acute myeloid leukemia (AML) is a phenotypically and genetically heterogeneous type of blood cancer, characterized by clonal expansion and loss of differentiation ability of immature myeloid progenitors. Clonal hematopoiesis promotes the outcompetition of normal cells by mutated hematopoietic stem cells in the bone marrow and peripheral blood. (1). AML is the most common type of acute leukemia (comprising 80% of all leukemic cases in adults and 15-20% in children) with an incidence rate of 3-5 cases per 100,000 population in the United States (2). With the currently approved therapeutic protocols, AML is cured in 35-40% of patients under 60 years old, and in 5-15% of patients over 60 years old, with a median survival of 5-10 months (3, 4). The development of resistance is the major mechanism behind the patients’ relapse. Identification of novel target genes and mechanisms that selectively control AML cell growth and disease progression may facilitate the development of new treatment approaches that specifically target the AML cells with less harmful effects on normal cells.

In AML genomes, transcription factors (TFs) are among the most frequently mutated and dysregulated genes, including chromosomal translocations in CBF, RARα, RUNX1 and the HOX gene family, but also point mutations targeting, for example, C/EBPα, P53 and RUNX1, are common aberration in AML genomes (5,6). TFs and their coactivators were early recognized for their involvement in AML development as oncogenic drivers (7). Thus, in order to improve the clinical outcome for patients with AML, it is important to gain further understanding of the molecular mechanisms promoting AML pathogenesis and progression governed by TFs.

GTF2IRD1 belongs to the TF II-I family and was first discovered in yeast (8). It is characterized by multiple helix-loop-helix-like I-repeat domains (9). It can function as either an activator or repressor depending on the cell type, target gene and its context (9–11). GTF2IRD1 is perhaps best known for its implicated role in Williams-Beuren syndrome, where GTF2IRD1 and the homolog GTF2I are part of a microdeletion, causing symptoms such as cardiovascular disease, a distinctive craniofacial appearance, intellectual disability and hypersociability (12). In addition, GTF2IRD1 has been reported to have an important role in different types of cancer, including colorectal cancer (13–15), pancreatic cancer (16) and breast tumor formation (17,18). However, the role of GT2IRD1 in normal and malignant hematopoiesis is largely unknown.

In this study, we used shRNA-based RNA interference (RNAi) screens at a large scale to identify novel TFs that were required for cell growth of *MLL-AF9* transformed AML cells. We discovered GTF2IRD1 as being critical for AML cell growth but not primary normal hematopoietic cells. In syngeneic and patient derived xenograft mouse models for AML, inhibition of GTF2IRD1 significantly delayed disease onset and prevented maintenance of patient-derived leukemia-initiating cells (LICs). Furthermore, we have shown that GTF2IRD1 functions as a transcriptional repressor and inhibition of the TF resulted in accumulation of quiescent AML cells, which was consistent with downregulation of E2F targets, revealed by RNA sequencing (RNA-seq) analysis.

## Results

### Identification of transcription factors required for cell growth of AML cells, using large-scale shRNA-based RNAi screens

To identify novel TFs that are required for cell growth of AML cells carrying the *MLL-AF9* translocation, we performed shRNA-based screens on the NOMO-1, THP-1 and genetically defined mouse AML cells, all carrying the MLL-AF9 fusion oncogene. The barcoded lentiviral RNAi library consisted of 27,500 shRNAs targeting approximately 5,000 annotated genes enriched for signaling pathways and the majority of annotated TFs (Cellecta Inc.). The AML cells were transduced by viral transfer in a pooled format as described in Figure 1A and the cells were harvested for genomic DNA extraction at an initial time point and after ten cell divisions. Effects on cell growth of the shRNA-targeted genes were determined by next-generation sequencing of PCR-amplified barcodes, originating from the genomic DNA of the AML cells (Supplementary table 1). By calculating the ratio of the isolated barcoded shRNAs at the initial time point with ten cell divisions at the population level, 647 target genes with at least a five-fold reduction were overlapping between all three cell lines used in the screens (Figure 1B, Supplementary Table 1). With a focus on TFs, 38 of the total 647 hits represented annotated TFs (Figure 1C). Among the identified 38 TFs, several have previously been reported to have a role in AML, including MEIS1, IKZF1/2, EZH1, HES1 and MYC, suggesting the validity of the screens, whereas 10 of the 31 TFs have not been reported to have a role in AML (Figure 1D). GTF2IRD1 was one of the TFs with the highest effects in reduced cell proliferation upon inhibition. Due to its remarkable overexpression in subpopulations enriched for leukemia-initiating cells compared to normal counterpart cells (19), and its important role in different cancer types (13–18), we decided to focus on GTF2IRD1 to validate its functional effects and the underlying mechanism.

**Figure 1.**
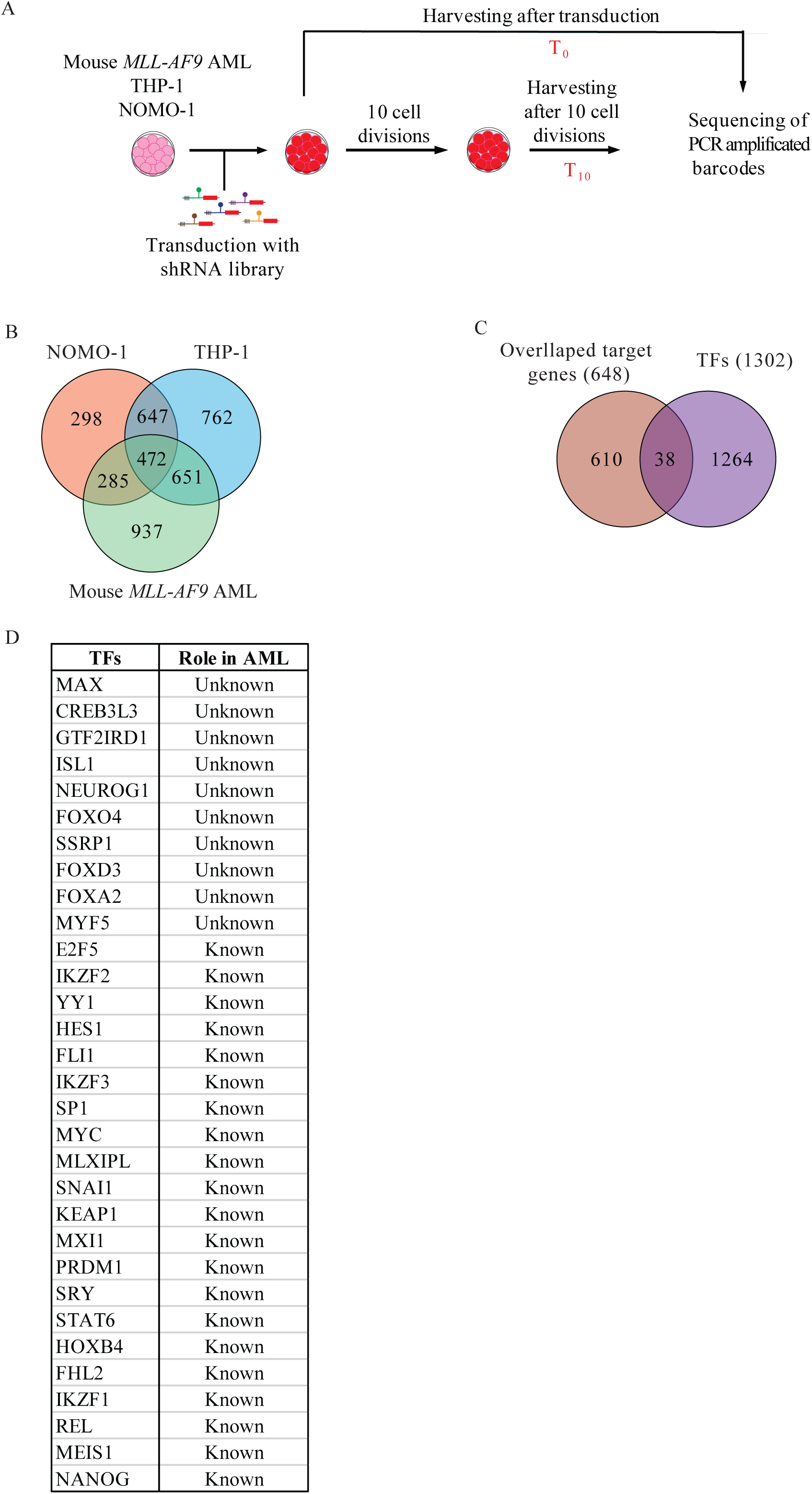
Largescale RNAi screens identified GTF2IRD1 as a novel transcription factor with a potential role in AML cell growth. **A**. Experimental scheme for identification of target genes by performing large-scale shRNA screens in AML cells. **B**. Venn diagram showing overlapping target genes between mouse *MLL-AF9* AML and human THP-1 and NOMO-1 AML cell lines that caused >five-fold reduction in expansion of the cells in the screens after ten cells divisions. **C.** Venn diagram showing overlap between the 648 identified target genes from the screens, to all annotated TFs (Pfam database). **D.** The table represent the 38 known and unknown TFs identified from the screens.

### Inhibition of *GTF2IRD1* prevents expansion of AML cell lines and delays disease onset in a mouse model for AML

To validate the findings from the screen, we performed cell growth competition assays by first co-culturing mouse *MLL-AF9* AML cells transduced with an shRNA vector targeting *Gtf2ird1* co-expressing RFP reporter, with control cells expressing a negative scrambled shRNA and GFP reporter. Efficient inhibition of *Gtf2ird1* mRNA in mouse *MLL-AF9* AML cells (Figure 2A), and flow cytometry analysis revealed a marked reduction in the number of live AML *Gtf2ird1*-inhibited cells, as compared to the mock control cells (Figure 2B and C). Likewise, a significant reduction in mRNA levels upon inhibition of *GTF2IRD1* in the human THP-1 and NOMO-1 AML cells expressing the *MLL-AF9* translocation (Figure 2D), prevented cell growth of both cell lines (Figure 2E and F), indicating that *GTF2IRD1* is essential for different AML cells carrying the *MLL-AF9* translocation. Similarly, inhibition of *GTF2IRD1* in non-*MLL*-rearranged leukemic cells, including HL-60, K-562 or NB-4 cells (Figure 2G), significantly reduced cell growth of these cells *in vitro,* showing that *GTF2IRD1* is also important for AML cells not carrying the *MLL-AF9* translocation (Figure 2H and I).

**Figure 2.**
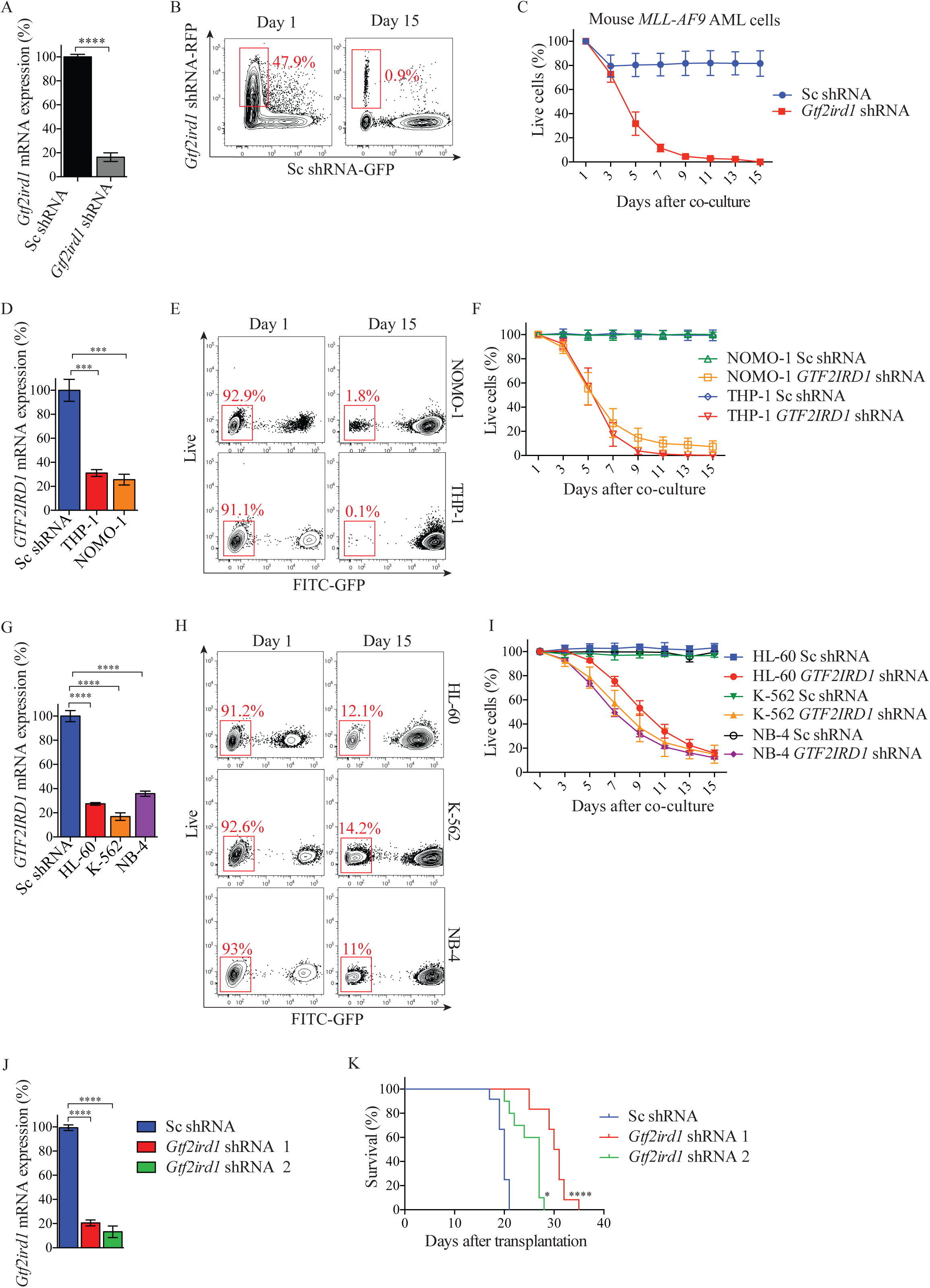
GTF2IRD1 inhibition impairs mouse and human leukemia cell growth and prolongs survival in mice with AML. **A.** Real time qPCR analysis of mRNA levels of *Gtf2ird1* assessed in mouse AML cells after transduction with vectors targeting *Gtf2ird1* or control (Sc) shRNAs. The mRNA levels were normalized to *Gapdh*. The data is presented as mean ±S.E.M., ****p<0.001, (unpaired *t*-test). **B.** Representative flow cytometric analysis of the cell growth competition assay in (C), where the mouse AML cells transduced with lentiviral vectors expressing shRNA against *Gtf2ird1* and RFP, or negative control vectors expressing scrambled shRNA and GFP (Sc), were mixed and propagated for 15 days. **C**. Line chart of the cell growth competition assay where the relative ratio of the RFP^+^ *Gtf2ird1* inhibited cells and the control cells was dynamically monitored by flow cytometric analysis. Percentages of live RFP^+^ cells transduced with *Gtf2ird1* shRNA (red line) were normalized to 100% based on the Day 1 numbers. **D**. Real time qPCR analysis of mRNA levels of *GTF2IRD1* in THP-1 and NOMO-1 cells after transduction with vectors targeting *GTF2IRD1* or control (Sc) shRNAs, (normalized to UBC) in cells transduced with Sc control (black bar) or GTF2IRD1 shRNA (grey bar) as indicated. The data is presented as mean ±S.E.M., ****p<0.001, (unpaired *t*-test). **E**. Representative flow cytometric analysis of THP-1 cells transduced with lentiviral vectors expressing shRNA against *GTF2IRD1*, or control negative control vectors expressing scrambled shRNA and GFP (Sc). The transduced cells were used in the cell growth competition assay in (F) and propagated for 15 days **F**. Line chart showing the relative ratio of the live cells and the control GFP^+^ cells dynamically monitored by flow cytometric analysis, at the indicated time points for 15 days after mixing the cells. Percentages of live RFP^+^ cells transduced with *GTF2IRD1* shRNA were normalized to 100% based on the Day 1 values and shown in bar graphs compared to Sc control shRNA. **G**. Real time qPCR analysis of mRNA levels of *GTF2IRD1* in HL-60, K562 and NB4 cells after transduction with vectors targeting *GTF2IRD1* or control (Sc) shRNAs, (normalized to UBC) in cells transduced with Sc control (black bar) or GTF2IRD1 shRNA (grey bar) as indicated. The data is presented as mean ±S.E.M., ****p<0.001, (unpaired *t*-test). **H**. Representative flow cytometric analysis of HL-60, K-562 and NB-4 cells, cells transduced with lentiviral vectors expressing shRNA against *GTF2IRD1*, or control negative control vectors expressing scrambled shRNA and GFP (Sc). The transduced cells were monitored as described in (E). **I**. Line chart showing the relative ratio of the live cells and the control GFP^+^ cells dynamically monitored by flow cytometric analysis, at the indicated time points for 15 days after mixing the cells. Percentages of live RFP^+^ cells transduced with *GTF2IRD1* shRNA were normalized to 100% based on the Day 1 values and shown in bar graphs compared to Sc control shRNA. **J.** Real time analysis of *Gtf2ird1* mRNA level (normalized to Gapdh) in *MLL-AF9* derived AML cells transduced with control (Sc) or two independent *Gtf2ird1* shRNAs before transplanting into sublethally radiated mice. **K**. Kaplan-Meier survival curves are shown for mice transplanted with mouse *MLL-AF9* AML cells (200,000 cells/mouse), transduced with shRNA against *Gtf2ird1* or negative control vectors. n = 10 in each cohort. The data is presented as mean ±S.E.M., *p<0.05, ***p<0.005, ****p<0.001, (unpaired t-test).

To investigate the role of *Gtf2ird1* in AML disease progression, we transplanted mouse *MLL-AF9* AML cells transduced with two individual *Gtf2ird1* shRNAs or a negative scramble control shRNA into recipient mice. Inhibition of *Gtf2ird1* was robust as compared to the control (Figure 2J). Mice that received control cells (Sc shRNA) develop AML within 18-21 days after transplantation (Figure 2J). However, in sharp contrast, mice that received AML cells transduced with the two individual shRNAs survived significantly longer than the control group (Figure 2K), showing that inhibition of *Gtf2ird1* delays disease onset in an immunocompetent transplantation mouse model for AML.

### *GTF2IRD1* is not essential for normal hematopoietic cell growth and function

We then asked whether inhibition of *GTF2IRD1* affects the expansion and differentiation of normal mouse hematopoietic stem and progenitor cells (HSPCs). Dynamic flow cytometric analysis of the HSPCs using an *in vitro* cell growth competition assay, showed no significant difference in expansion between *Gtf2ird1*-inhibited cells as compared to control cells transduced with a non-targeting scramble vector, during 7 days of culturing (Figure 3A-C). Furthermore, neither the total number of colonies nor mixed myeloid-erythroid CFUs (CFU-GEM) nor granulocyte/macrophage CFUs (CFU-GM) were significantly affected in terms of colony numbers from cells transduced with shRNA against *Gtf2ird1* as compared to cells carrying a non-targeting negative control vector (Figure 3D and E). This indicated that *Gtf2ird1* was not essential for proliferation, survival or differentiation of mouse HSPCs *in vitro*.

**Figure 3.**
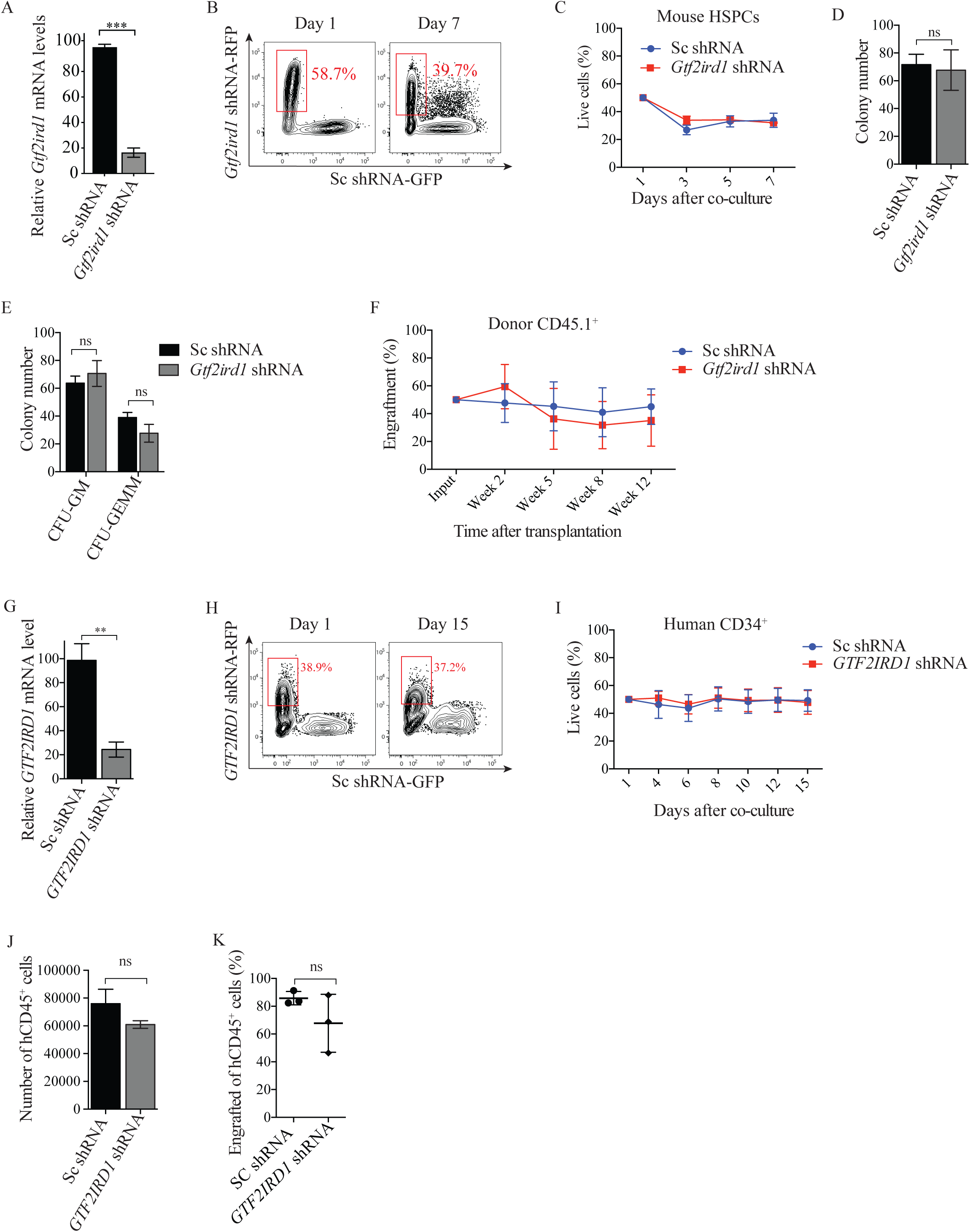
*GTF2IRD1* suppression is not essential for cell growth of normal hematopoietic cells *in vitro* and *in vivo*. **A.** Real time qPCR analysis of mRNA levels of *Gtf2ird1* assessed in mouse HSPCs after transduction with vectors targeting *Gtf2ird1* or control (Sc) shRNAs. The mRNA levels were normalized to *Gapdh*. The data is presented as mean ±S.E.M., ***p<0.005, (unpaired t-test). **B.** Representative flow cytometry analysis charts showing percentage of live RFP^+^ *Gtf2ird1* inhibited mouse HSPCs (*Gtf2ird1* shRNA-RFP) relative to GFP^+^ control cells (Sc shRNA-GFP) at the indicated days after mixing the cells for the *in vitro* growth competition assay in (C). **C**. Line charts depicting the percentage of live RFP^+^ cells normalized to 100% with GFP^+^ cells after mixing the HSPCs, determined by flow cytometry analysis at the indicated time points. **D** and **E**. Clonogenic progenitor assays of sorted *Gtf2ird1* inhibited or control HSPCs where equal cell numbers were plated in in methylcellulose and total colonies were counted after seven days CFU-GM and CFU-GEMM (D), or after ten days (E), of clonal growth. Counts of different colony types are shown as mean ±SEM of n = 3, ns = non-significant, (unpaired t-test). **F**. Line charts showing the percentage of engraftment in the BM of recipient mouse after transplantation of sorted *Gtf2ird1* RFP^+^ inhibited CD45.1 and GFP^+^ control cells into a congenic transplantation model. Live RFP^+^ cells were normalized to 100% after mixing the HSPCs, determined by flow cytometry analysis at after 2, 5, 8 and 12 weeks post transplantation for total donor CD45.1^+^ cells. The data is represented as the mean ±SEM, n=6. **G**. Real time qPCR analysis of mRNA levels of *GTF2IRD1* in CD34^+^ enriched cells after transduction with vectors targeting *GTF2IRD1* (*GTF2IRD1* shRNA, grey bar) or control shRNAs, (Sc control (black bar), normalized to UBC. The data is presented as mean ±S.E.M., **p<0.01, (unpaired t-test). **H**. Representative flow cytometry plots of percentage of live RFP^+^ *GTF2IRD1* inhibited human HSPCs (*GTF2IRD1* shRNA-RFP) relative to GFP^+^ control cells (Sc shRNA-GFP) at the indicated days after mixing the cells for the *in vitro* growth competition assay in (I). **I**. Line chart of cell growth competition assay where CD34^+^ CB cells transduced with shRNAs against *GTF2IRD1* were mixed and expanded in suspension with supplemented cytokines for 15 days. The number of viable cells were determined by flow cytometric analysis and the data is shown as mean ±SEM of n = 3, ns = non-significant (unpaired *t*-test). **J.** Quantification of long termed expanded live normal CD34^+^ UCBs transduced with *GTF2IRD1* targeting vectors or negative scramble control vectors after culturing the cells in suspension for 3 weeks. The number of human CD45^+^ UCBs was determined by flow cytometric analysis after three weeks of culturing. The data is represented as mean ±SEM of n = 3, ns = non-significant, (unpaired *t*-test). **K.** Box plot showing the percentage of live CD45^+^ UCBs transduced with *GTF2IRD1* targeting shRNA or scramble control vector, eight weeks after transplantation of 50,000 cells into humanized NSG-SGM3 recipient mice by intra femural injection. The BM of the xenografted mice was analyzed for human CD45^+^ engraftment after transplantation by flow cytometry. Results are shown as mean ±SEM of n = 3 mice per experiment, ns = non-significant, (unpaired t-test).

Next, we investigated the importance of *Gtf2ird1* in engraftment and performed intravenous transplantation of sorted ckit^+^ cells transduced with *Gtf2ird1* or a non-targeting negative control vector into lethally irradiated recipient mice. Flow cytometric analysis of BMs from the transplanted mice revealed minor changes in the engraftment of mice transplanted with *Gtf2ird1*-inhibited cells compared to the control group, suggesting that *Gtf2ird1* is not critical for engraftment of HSCs in the bone marrow (Figure 3F).

To investigate the importance of *GTF2IRD1* in normal human hematopoietic cell proliferation and survival, we tested the effects on cell growth of umbilical cord blood cells (UCBs) transduced with shRNA vectors against *GTF2IRD1* compared to co-cultured control cells (Figure 3G). Dynamic flow cytometric analysis of the mixed cells showed non-significant differences in expansion in suspension of the UCBs transduced with an shRNA against GTF2IRD1 compared to the control cells (Supplementary Table 1, Figure 3H and I). To explore the importance of *GTF2IRD1* in primary human hematopoietic cells in a more long-term perspective, we again transduced UCBs with *GTF2IRD1-specific* shRNAs and control vectors, and maintained the UCBs in a competitive setting on stromal cell feeder layers for 3 weeks. Flow cytometric analysis showed that the number of live CD45-positive (CD45^+^) UCBs was not significantly changed between the *GTF2IRD1*-targeted cells and control cells (Figure 3J). Consistent with the transplantation studies in the syngeneic mice in Figure 3F, intravenous tail injections of sorted human CD34-positive (CD34^+^) UCBs into humanized NSG-SGM3 recipient mice showed non-significant differences in engraftment between the *GTF2IRD1*-targeted cells compared to the control cells (Figure 3K). Together, these data suggested that *GTF2IRD1* is not essential for normal hematopoietic viability and differentiation *in vitro* as well as *in vivo*.

### CRISPR-Cas9-mediated disruption of *GFT2IRD1* prevents the expansion of AML cells

To rule out possible off-target effects associated with the shRNA-mediated loss of function experiments, we conducted CRISPR-Cas9-mediated gene editing experiments in combination with cell growth competition assays. THP-1 and MV4-11 AML cells, which constitutively expressed the Cas9 nuclease, were transduced with a guide RNA (gRNA) vector homologous to the coding region of human *GTF2IRD1* (Figure 4A). The *GTF2IRD1* gRNA co-expressing the fluorescent marker (iRFP670) caused a significant disruption of *GTF2IRD1*, using a non-targeting control gRNA expressing BFP (Figure 4B). By co-culturing the cells and monitoring the relative ratio of the cells with flow cytometry in the so-called competition assay, we observed a strong reduction of live cells transduced with *GTF2IRD1* gRNA compared to control cells, in both THP-1 and MV4-11 AML cells as compared to the control cells (Figure 4C, D, E).

**Figure 4.**
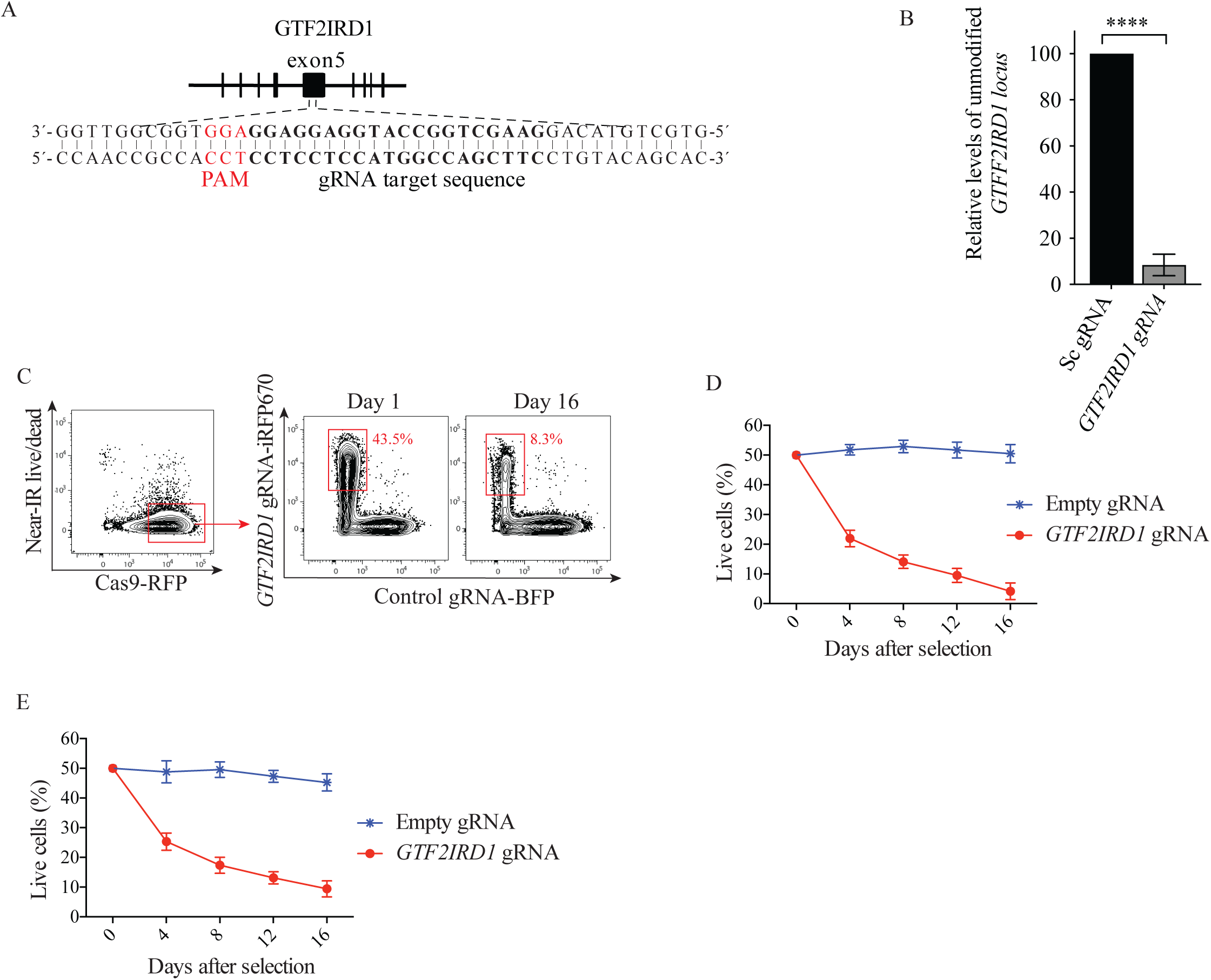
CRISPR-Cas9 gene editing confirmed that *GTF2IRD1* is essential for expansion of human leukemic cells. **A.** Schematic image of gRNA design and targeting location at exon 5 of *GTF2IRD1*. The gRNA target sequence is highlighted in bold letters and the PAM sequence is highlighted in blue. **B**. Bar diagram showing the level of gene disruption 72 hours after transduction of THP-1 cells, for the gRNA used in C-E, determined by digital PCR analysis. The data is shown as mean ±SEM of n = 3, (unpaired *t*-test) ****p<0.001. **C.** Representative flow cytometry analysis of percentage of THP-1 AML cells stably expressing the Cas9 nuclease and the mCherry reporter transduced with a gRNA vector expressing gRNA against *GTF2IRD1* (iRFP-670^+^) or control gRNA (BFP^+^), at the day of mixing the cells and after 16 days of co-culturing for the cell growth competition assay shown in (C). **D** and **E**. Line chart of cell growth competition assay showing the relative abundance of *GTF2IRD1* (iRFP-670^+^) THP-1 cells (C) or NOMO-1 cells (D), or control gRNA (BFP^+^) cells, determined by flow cytometric analysis at the indicated time points. The data is shown as mean ±SEM of n = 3, ns = non-significant (unpaired *t*-test).

### Inhibition of *GTF2IRD1* reduces the frequency of primary adult and childhood AML cells, including LICs *ex vivo*

Having demonstrated a selective importance of *GTF2IRD1* in various AML cell lines and disease progression in animal models for AML, we wanted to investigate the role of *GTF2IRD1* in primary adult and childhood AML samples, including the LIC population. For that purpose, the patient samples from both adults and children were transduced with either *GTF2IRD1* targeting vectors or negative control vectors (used in Figure 2D-H) and maintained in a co-culture system with stromal cells (Supplementary Table 2 staining panel for sorting and Supplementary Table 3 AML patients’ samples annotation). Although one of the samples (”Adult AML 7”) did not show a significant difference due to high variation of the cells in the negative control sample, flow cytometric analysis of *GTF2IRD1* targeted cells caused a consistent reduction in maintenance of primary CD45^+^ AML bulk cells isolated from adult patients, compared to the cells transduced with a control vector (Figure 5A). Similarly, the Lin^−^ CD34^+^CD38^−^ subpopulation enriched for LICs, revealed a significant reduction in the frequency of cells upon inhibition of *GTF2IRD1* except for one sample (“Adult AML 5”), where the variation in the negative control cells was too large to show a statistically significant difference (Figure 5B). To investigate if the importance of *GTF2IRD1* was conserved in primary childhood AML cells, we conducted shRNA-based inhibition of *GTF2IRD1* in two childhood AML samples as described in Figure 5A and B. Consistent with the AML samples from adults, inhibition of *GTF2IRD1* significantly reduced the numbers of CD45^+^ bulk cells as compared to the control cells Figure 5C. We also observed a reduction of Lin^−^CD34^+^CD38^−^ childhood AML cells for one of the samples upon inhibition of *GTF2IRD1* (“Childhood AML 1”), whereas the reduction of cells in the second sample (“Childhood AML 2”) was not statistically significant (Figure 5D).

**Figure 5.**
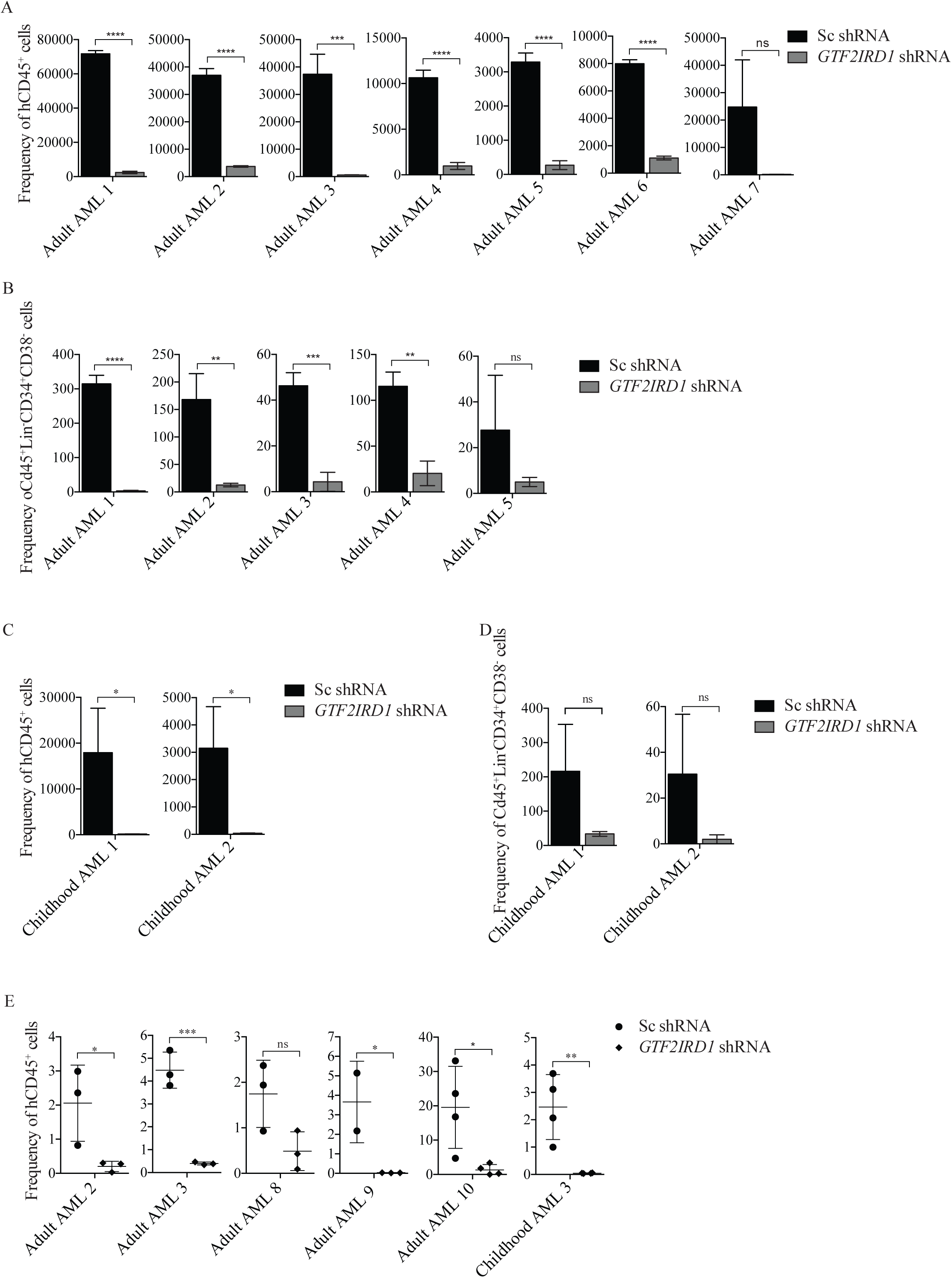
Inhibition of *GTF2IRD1* significantly reduce the frequency of primary adult and childhood AML cells, *ex vivo* and in a xenograft mouse model for AML. **A** and **B.** Bar diagram showing the frequency of patient derived CD45^+^ bulk cells form seven adult AML patient samples (A), or Lin^−^CD34^+^CD38^−^ enriched LICs from five adult AML patient samples, transduced with *GTF2IRD1* shRNA (grey bars) compared to Sc shRNA (black bars). AML patient samples were cultured in long term on MS-5 stromal cells. The data is shown as mean ±SEM of n = 3, ns = non-significant, **p<0.01, ***p<0.005, ****p<0.001 (unpaired t-test). **C** and **D**. Bar diagram of the absolute number of patient derived CD45^+^ bulk cells form two childhood AML patient samples (C), or Lin^−^CD34^+^CD38^−^ enriched LICs from the two childhood AML patient samples, transduced with *GTF2IRD1* shRNA (grey bars) compared to Sc shRNA (black bars). AML patient samples were cultured in long term on MS5 stromal cells. The data is shown as mean ±SEM of n = 3, ns = non-significant, *p<0.05. **E**. Box plot of the frequency of engrafted primary AML patient CD45^+^ cells transduced with GTF2IRD1 or Sc shRNA, isolated from the BM of recipient NSG-SGM3 mice after eight weeks. The frequencies of the engrafted cells were determined by flow cytometric analysis and each individual dot represents level of engraftment in an individual mouse. *p<0.05, **p<0.01, ***p<0.005 (unpaired *t*-test).

Importantly, intrafemoral injection of five adult and one childhood primary AML samples, with and without shRNA-mediated inhibition of *GTF2IRD1* into humanized NSG-SGM3 mice, except for one sample (“Adult AML 8“), showed a significant decrease in the total number of human CD45^+^ cells in the BM of the recipient mice as compared to the control cells. Together, these studies suggested that *GTF2IRD1* is essential for maintenance of both primary AML cells from both adults and children, including the LICs and for engraftment in mice.

### Downregulation of *Gtf2ird1* as a transcriptional repressor blocks AML cells in the G0 phase of the cell cycle without causing apoptosis

Next, we asked whether the observed essential role for *Gtf2ird1* in AML cells was associated with a function in cell cycle progression or cell apoptosis. Analysis of mouse *MLL-AF9* AML cells 72 hours after *Gtf2ird1* inhibition with significant mRNA reduction (Figure 6A) showed a strong accumulation of cells in the G0 phase of the cell cycle and a subsequent reduction in G1, S and G2-M phases, in comparison to the negative control cells (Figure 6B). In contrast, the percentage of early and late number of apoptotic cells was relatively low (between 1-3%) (Figure 6C), suggesting that the main cellular consequence to inhibition of *Gtf2ird1* was the accumulation of cells in the G0 phase of the cell cycle.

**Figure 6.**
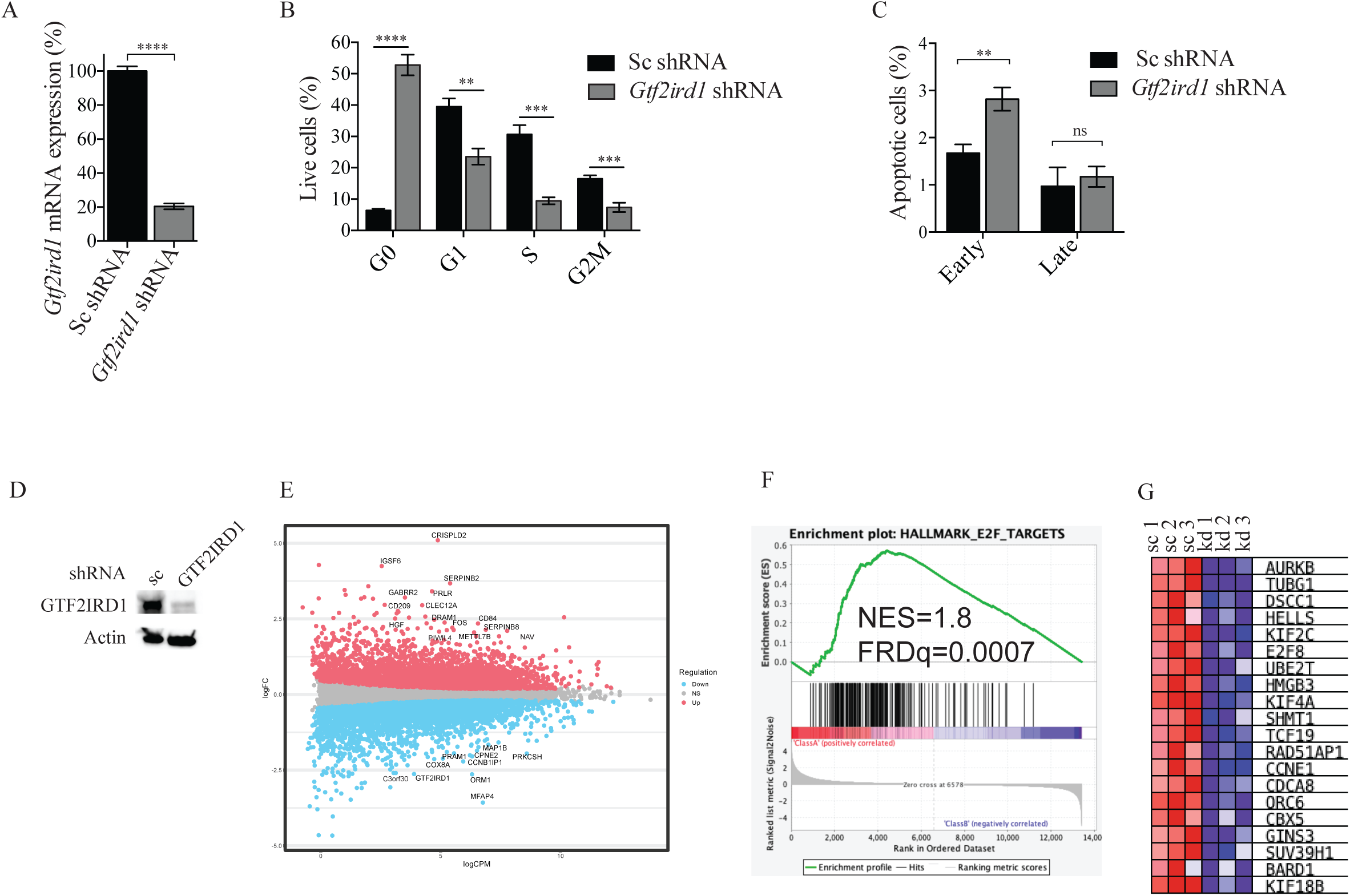
inhibition of GTF2IRD1 arrest AML cells in the G0 phase of the cell cycle and downregulates E2F targets. **A**. Real time qPCR analysis of mRNA levels of *Gtf2ird1* assessed in mouse *MLL-AF9* AML cells after transduction with vectors targeting *Gtf2ird1* or control (Sc) shRNAs. The mRNA levels were normalized to *Gapdh*. The data is presented as mean ±S.E.M., ***p<0.005, (unpaired *t*-test). **B**. Bar graphs illustrating flow cytometric quantification of mouse *MLL-AF9* AML cells transduced with *Gtf2ird1* shRNAs, or a negative control vector, stained with Ki-67 and DAPI, at 72 hours after transduction. The data is presented as mean ±S.E.M., n=3, **p<0.01, ***p<0.005, ****p<0.001 (unpaired *t*-test). **C**. mouse *MLL-AF9* AML cells transduced with *Gtf2ird1* shRNA, or a negative control vector, stained for apoptosis detection, at 72 hours after transduction. The data is presented as mean ±S.E.M., ns = non-significant,**p<0.01, (unpaired t-test), n=3. **D**. Western blot of protein extract from THP1 cells with shRNA-based inhibition of GTFF2IRD1 and control cells transduced with a negative control vector (sc), that were used for the RNA-seq analysis in Figure E-G. **E**. MA plot of RNA-seq analysis. Indicated genes represent top 20 leading edge genes **F**. GSEA enrichment plot of hallmark genes, showing the significant correlation to E2F targets upon GTFF2IRD1 inhibition in THP1 cells. NES score and FDRq scores are indicated in the diagram. **G**. Leading edge analysis of top 20 most significant genes affected as a consequence of biological triplicate experiments of GTF2IRD1 inhibition in THP1 cells.

### RNA profile pattern under *GTF2IRD1* suppression shows a correlation to E2F targets

To understand the molecular mechanism by which *GTFF2IRD1* was essential for AML cell growth, we performed differential gene expression profiling with RNA sequencing (RNA-seq) analysis with and without shRNA-mediated loss of function of THP-1 AML cells, using an shRNA against *GTFF2IRD1* and a negative control vector expressing a non-targeting shRNA (Supplementary Table 1, Figure 6D, Supplementary Fig. 1). Using statistical (FDR<0.1) and a two-fold change cutoff, 343 genes were found to be upregulated and 443 genes were downregulated as a consequence of inhibition of GTF2IRD1 (Figure 6E). To identify the dysregulated gene sets associated with the cellular response upon the *GTF2IRD1* inhibition, we performed gene set enrichment analysis (GSEA) (20, 21). This did not reveal any significantly altered pathways (KEGG, Biocarta, Wikipathways) or gene ontology. On the contrary, the expression changes in response to the inhibition of *GTFF2IRD1* compared to the hallmark gene sets were significantly correlated with the gene signature of E2F targets data sets (FDR Q-value=0.0007 and normalized enrichment score [NES]= 1.8.), (Figure 6F and G), suggesting that *GT2IRD1* is required for AML cell growth via regulation of E2F targets.

## Discussion

This study identifies *GTF2IRD1* as being required for cell growth of various AML cells *in vitro* and *in vivo*, including primary AML cells and LICs from adults and children, but not normal hematopoietic cells. The essential role of *GTFIRD1* in AML is associated with cell cycle inhibition at G0 phase.

Despite the genetic and epigenetic heterogeneity in primary AML samples (22), our results highlight a consistent importance of *GTF2IRD1* in the maintenance of primary childhood and adult AML cells with mixed genotypes *in vitro* and *in vivo,* including for enriched populations of LICs that drive the disease (23). Although GTF2IRD1 has previously not been reported to have a function in leukemia, it is overexpressed in AML hematopoietic stem and progenitor cells as compared to normal counterpart blood cells (19). In addition, there is a growing body of evidence that GTF2IRD1 has a functionally important role in additional cancer types, including colorectal cancer (13–15), pancreatic cancer (16), and breast cancer (17,18). It is therefore possible that GTF2IRD1 has a more general role in cancer than previously anticipated.

In the present study, we also show that the essential function of *GTF2IRD1* in AML is selective. Although further work is needed to address the underlying mechanisms of the observed leukemia-specific role, neither are normal primary hematopoietic cells significantly affected by functional inhibition of *GTF2IRD1,* nor is the reconstitution of the hematopoietic cells in animal models. Consistent with this, *Gtf2ird1* homozygote knockout mice are viable and display phenotypic features restricted to craniofacial abnormalities, cognitive defects and a mild growth deficiency (24, 25, 26). One possible explanation for the selective importance is the determined overexpression of GTF2IRD1 in subpopulations of AML cells as compared to normal counterpart cells (19).

In our study, inhibition of *Gtf2ird1* in AML cells results in a reduction of cell expansion via a mechanism that involves a cell cycle arrest and quiescent cells are resistant to loss of function of *Gtf2ird1*, manifested by an accumulation of the cells in the G0 phase of the cell cycle. Consistent with this, GSEA analysis revealed a significant correlation to downregulation of E2F targets upon inhibition of *GTF2IRD1*, and E2F effectors have a principal role in regulating the cell cycle of cells (27). The link of *GT2IRD1* to cell cycle regulation has also been reported for colon cancer (28).

Overall, we show for the first time that inhibition of GTF2IRD1 prevents disease progression and cell growth of primary AML cells but not normal hematopoietic cells. Chemical targeting of TFs via proteasome-mediated proteolysis is a remarkable approach for the treatment of various leukemia (29) and can be applied to a range of targets including newely identified by us GTF2IRD1. Thus, our findings might contribute to increased understanding of AML and provide therapeutic targets that may facilitate clinical solutions for the disease.

## Material and Methods

### Cell culture, cell lines and primary cells

The human AML cell lines; THP-1, NOMO-1, MV4-11, K-562, HL-60 and NB-4 (all purchased from DSMZ) were cultured in RPMI with 10% FBS, 100 U/mL penicillin and 100 ug/mL streptomycin at 37 °C, 5% CO2. The murine *MLL-AF9* AML cells (30) were cultured in the same conditions with 10 ng/mL rmIL-3 (R&D systems). Culturing of normal murine HSPCs and human CD34^+^ UCB cells has been described previously (31).

### Cell growth assays of primary AML samples

Isolation and culturing of the AML samples was carried out as previously explained (30). Briefly, transduced AML cells were cultured on MS-5 cells (DSMZ) in Myelocult media H5100 (StemCell Technologies Inc.) with rhIL-6, rhIL-3, rhFl3/Flk-2 ligand, rhTPO, and rhSCF and rhG-CSF (Stemcell technology), at a concentration of 20 ng/mL.

### Vectors, molecular cloning and knockout analysis

The inducible shRNA and gRNA vectors were generated as described before (30). gRNAs were designed by Optimized CRISPR Design - MIT (http://crispr.mit.edu) provided by the Zhang laboratory, and cloned into the inducible gRNA expression vectors. The constitutive lentiviral-based Cas9-mCherry vector was kindly provided by Marco Herold (31). CRISPR/Cas9 growth assay was explained in a previous article (30). All oligos used in the study are listed in Supplementary Table 1.

### Transfection and Transduction

Transfection and transduction were performed as previously described (30).

### Large-scale shRNA screens

Large scale shRNA screens were described in detail in a previous study (30). Briefly, around 3*×*10^8^ cells (THP-1, NOMO-1 and murine AML) were transduced with pooled barcoded lentiviral shRNA libraries targeting genes playing a role in signalling pathway (Module 1, Cellecta Inc). The transduction efficiency was kept below 25% to minimize the number of cells with multiple integrations of the vectors. Half of the cells were collected for an initial input control (T0), 48 hours after selection with puromycin (2 μg/mL) (Invitrogen) and the remaining cells were harvested 10 cell divisions. The barcodes were amplified from genomic DNA and the purified barcodes were used for next-generation sequencing (HiSeq 2000, Illumina).

The ratios of the barcodes from the two screens were calculated by first dividing the normalized abundance of each barcode after T10 by the abundance at the initial time point (T0) separately for each cell line.

### Flow cytometric analysis and sorting

Flow cytometry and sorting were performed as previously described (29). Primary AML cells were harvested and stained with human CD45, and cells were analyzed by a high-throughput automated plate reader (BD LSRFortessa). Dead cells were excluded using the Near-IR Live/Dead marker (Invitrogen). For the cell growth competition assays, cells washed with PBS were stained with Near-IR Live/Dead marker in a 96-well plate, and then a high-throughput automated plate reader was used to detect the absolute number of live cells.

To determine the level of engraftment of human AML cells, the isolated BMs were stained with human anti-CD45 and Near-IR Live/Dead marker to detect live cells. All FACS Analysis was performed by FlowJo Version 9.3.3 software (TreeStar). All antibodies used in the study are listed in Supplementary Table 3.

### RNA extraction and quantitative real-time PCR (qPCR)

RNA was extracted using RNeasy Mini Kit (Qiagen), following the manufacturer’s instructions. cDNA synthesis was performed with 500 ng of RNA by using a cDNA synthesis kit (Fermentas or BioRad). qPCR was performed using an ABI Thermal Cycler, and qPCR primers were purchased from Eurofins (primers are presented in Supplementary Table 1).

### AML mouse models

All the mice were kept in a pathogen-free animal facility in Karolinska Institutet, Huddinge, Sweden. The C57BL/6 wild type mice and the NOD-scid IL2Rgnull-3/GM/SF, NSG-SGM3 mice were purchased from The Jackson Laboratory. All of the transplanted mice were investigated daily for symptoms of leukemogenesis and disease progression was monitored by complete blood tests.

The mouse MLL-AF9 induced AML cells and AML mouse model was generated as previously described (30, 31). The generated MLL-AF9 AML cell line was used for further transplantation experiments or *in vitro* experiments.

For the Kaplan-Meier survival analysis, wild type CD45.2 C57BL/6 mice were used for transplantation of murine MLL-AF9 AML cells. 100,000 AML cells transduced with lentiviral shRNA vectors, selected with puromycin (2 μg/mL) and were intravenously transplanted into non-irradiated recipient mice. After disease onset, the mice were euthanized.

The experiments using humanized NSG-SGM3 mice were performed on animals aged six to ten weeks. Recipient mice were subjected to sub-lethal irradiation (220 cGy) and six to 12 hours post irradiation the primary childhood AML patient BM samples were transplanted via intra-femur injection using doses of 100,000-500,000 cells/mouse. BM samples were collected from the bones of euthanized primary recipient mice 6-8 weeks post transplantation and the primary leukemic cells were analyzed with flow cytometric analysis for donor cell engraftment (See Supplementary Table 2 for antibodies).

### Cell cycle and apoptosis analysis

The procedure of cell cycle assay and apoptosis assay were explained before (30).

### RNA-Sequencing

Total RNA was extracted from human AML THP-1 cells by miRNeasy Kit (Qiagen), 96 hours after transduction of the cells with shRNA against *GTFF2IRD1* and a negative control vector expressing a non-targeting shRNA and after 96 hours selection on puromycine (2 μg/mL). Ilumina NextSeq 2000 was used to evaluate DEGs (differentially expressed genes) and 7-20 million reads/sample were obtained. The process was described in detail in the previous study (30).

### Western Blotting

Total proteins were prepared from cells using RIPA Buffer with Protease Inhibitor Cocktail (sc24948, Santa Cruz Biotechnology) and sonicated at 4°C/ 5 min 30s ON/30s OFF. Protein quantified was performed using the Bradford assay (Bio-Rad). 250 ug protein was prepared in the 1x BioRad lammeli buffer with DTT (#1610747, BioRad). Proteins were separated in 10% TGX gels (Biorad) and transferred to HyBond ECL nitrocellulose membranes (GE Health Life Sciences) by wet transfer in 20% methanol.

Nitrocellulose membranes were blocked with 5% nonfat dried milk (170-6404, Biorad) with 1X PBS (D8537, Sigma) and 0.2% Tween 20 (P9416, Sigma) before incubating with the primary polyclonal antibodies GTF2IRD1 (ProteinTech 17052-1-AP) and A5441, Sigma beta-actin respectively. After incubating with secondary antibodies, the detection of proteins was performed by using enhanced chemiluminescence kit (32106, Thermo Scientific) and the image was captured by a LI-COR Odyssey Fc Imager system. Quantification of the relative intensity of the bands was performed by using Image Studio Lite (LI-COR Biosciences).

### Statistical analysis

Statistical analysis was done with GraphPad Prism version 6, and p-values of <0.05 were considered statistically significant. GSEA analysis was done using log2 transformed expression values and the Molecular Signature Data Base, MSigDB (http://software.broadinstitute.org/gsea/msigdb/annotate.jsp), applying “Compute Overlaps” tool and “Hallmark gene sets”, “Reactome gene sets”.

## Ethics Statement

All experiments were done with the ethical standards and according to the Declaration of Helsinki and according to national and international guidelines and has been approved by the authors’ institutional review board. The primary AML childhood BM samples were obtained from Uppsala biobank (reg.nr 827) after approval from NOPHO - Nordic Society of Pediatric Hematolgy and Oncology and the local ethical committee at Stockholm (ethical permit: 2017-1059-32). All animal experiments were approved by the local ethics committee in Linköping, Sweden and the Swedish board of agriculture (Ethical permit number; ID 530). UCB samples were obtained from the Karolinska Hospital (Stockholm, Sweden) with informed consent of the parents.

## Acknowledgements

We thank Tim Somervaille for kindly providing the MLL-AF9 vector and Marco Herold for kindly providing the Cas9-mCherry vector. We also would like to thank NOPHO and Josefine Palle for providing the primary childhood AML cells as well as Cecilia Götherström and the National Cord Blood bank at Karolinska University Hospital for providing the UCBs. We would like to acknowledge the MedH Core Flow Cytometry facility (Karolinska Institutet), supported by KI/SLL, for providing cell sorting services, cell analysis services, technical expertise and scientific input. We also would like to thank the Affymetrix core facility at Neo, BEA, Bioinformatics and Expression Analysis, which is supported by the board of research at the Karolinska Institute and the research committee at the Karolinska University Hospital.

## Author contributions

All authors have read and provided input into the manuscript. Y.H. designed and performed most experiments, analyzed data and wrote the manuscript. G.T. deisgned and performed experiments, animal work, cell growth assays and RNA sequencing preparation. JZ. performed the RNA-seq experiments, provided input into the design of the project. N.K. M.D. provided input into the study and helped with long-term expansion of hematopoietic cells and flow cytometric analysis. A.H. performed analysis of RNA-Seq data and the shRNA screens. J.B. cloned and designed the inducible gRNA and shRNA vectors. H.C. helped with the RNA-seq analysis. M.A. designed the gRNA and shRNA vectors and provided input in the project. H.Q. helped to design the studies, drafting the manuscript, analysis of the experiment and supervision of Y.H. J.W. Conceived the project, supervised the study, analyzed the data, interpreted result and wrote the manuscript. All authors participated in editing of the manuscript.

