## Supplemental material for "Identification of GTF2IRD1 as a novel transcription factor essential for acute myeloid leukemia"

**Supplemental Table 1.** All oligos and gRNAs used in this study.

| Target site | Application | Forward oligo (5′>3′) | Reverse oligo (5′>3′) |
|---|---|---|---|
| m*Gtf2ird1* | qPCR | TCGGATGTGTACCTGCTGCA | TCCCTGAGCAGCCTCTCATA |
| h*GTF2IRD1* | qPCR | CCAAAGACACCACGAAGCTGGA | CCACAGTCTTCAGACATGCTG C |
| h*UBC* | qPCR | CTGGAAGATGGTCGTACCCTG | GGTCTTGCCAGTGAGTGTCT |
| m*Hprt1* | qPCR | GAAAAGGACCTCTCGAAGTGTTG | CACTAATGACACAAACGTGAT TCAAA |
| dHTS | 1st PCR barcodes | TCTCTGGCAAGCAAAAGACGGC ATA | TGCCATTTGTCTCGAGGTCGA GAA |
| dGex | 2nd PCR barcodes | CAAGCAGAAGACGGCATACGAG A | AATGATACGGCGACCACCGA GA |
| GexSeqN | HT seq. barcodes | ACAGTCCGAAACCCCAAACGCA CGAA | |

**Supplemental Table 2.** All antibodies used in this study.

| Antibody | Conjugated | Clone | Source |
|---|---|---|---|
| Anti-mouse TER-119 | Purified | TER-119 | Biolegend |
| Anti-mouse CD3 | Purified | 17A2 | Biolegend |
| Anti-mouse/Human CD45R/B220 | Purified | RA3-6B2 | Biolegend |
| Anti-mouse Ly-6G/Ly-6C (Gr-1) | Purified | RB6-8C5 | Biolegend |
| Anti-mouse NK1.1 | Purified | PK136 | Biolegend |
| Anti-mouse CD11B | Purified | M1/70 | Biolegend |

| | | | |
|---|---|---|---|
| Anti-mouse CD16/32 | Purified | 93 | Biolegend |
| Anti-mouse CD117 (c-kit) | APC-Cy7 | 2B8 | Biolegend |
| Ki-67 | FITC | B56 | BD |
| Anti-human CD20 | PE-Cy5 | 2H7 | Biolegend |
| Anti-human CD4 | PE-Cy5 | RPA-T4 | Biolegend |
| Anti-human CD8 | PE-Cy5 | RPA-T8 | Biolegend |
| Anti-human CD2 | PE-Cy5 | RPA-2.10 | Biolegend |
| Anti-human CD56 | PE-Cy5 | HCD56 | Biolegend |
| Anti-human CD235a | PE-Cy5 | HIR2 | Biolegend |
| Anti-human CD3 | PE-Cy5 | HIT3a | Biolegend |
| Anti-human CD19 | PE-Cy5 | HIB19 | Biolegend |
| Anti-human CD34 | APC | 581 | Biolegend |
| Anti-human CD38 | PE-Cy7 | HB7 | BD |
| Anti-human CD45 | BV786 | HI30 | BD |

**Supplemental Table 3.** Primary AML patient samples related to Figure 5.

| Sample | Type | ID | MLL Translocation | Experiment | Figure |
|---|---|---|---|---|---|
| AML 1 | Adult | 2009-022 | Non MLL | *Ex vivo* | 5A, B |
| AML 2 | Adult | 2011-031 | Non MLL | Ex vivo and *in vivo* | 5A, B and E |

| | | | | | |
|---|---|---|---|---|---|
| AML 3 | Adult | 2011-055 | Non MLL | *Ex vivo* and *in vivo* | 5A, B and E |
| AML 4 | Adult | 2008-23 | Non MLL | *Ex vivo* | 5A, B |
| AML 5 | Adult | 2008-27 | Non MLL | *Ex vivo* | 5A, B |
| AML 6 | Adult | 2012-22 | Non MLL | *Ex vivo* | 5A |
| AML 7 | Adult | 2012-14 | Non MLL | *Ex vivo* | 5A |
| AML 8 | Adult | 2011-48 | Non MLL | *In vivo* | 5E |
| AML 9 | Adult | 2012-24 | Non MLL | *In vivo* | 5E |
| AML 10 | Adult | 2014-16 | Non MLL | *In vivo* | 5E |
| AML 1 | Childhood | 03-215 | Non MLL | *Ex vivo* | 5C and D |
| AML 2 | Childhood | 06-013 | Non MLL | *Ex vivo* | 5C and D |
| AML 3 | Childhood | 03-316 | Non MLL | *In vivo* | 5E |

**Supplementary Figure 1**

Real time qPCR analysis of mRNA levels of *GTF2IRD1* assessed in human THP-1 AML cells after transduction with vectors targeting *GTF2IRD1* or control (Sc) shRNAs. The mRNA levels were normalized to *GAPDH*.

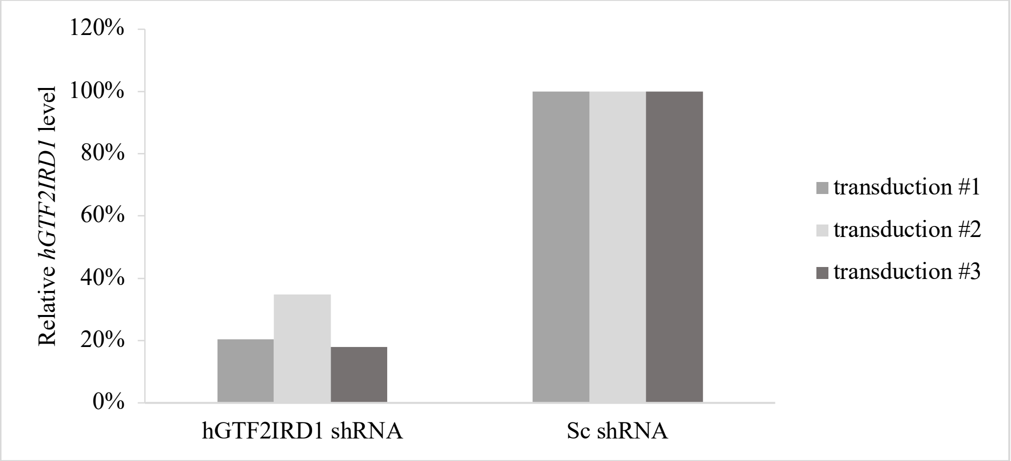
